# Polyphosphate and pyoverdine synthesis modulates *Pseudomonas aeruginosa* PAO1 virulence in a zebrafish model

**DOI:** 10.64898/2026.03.19.712776

**Authors:** Javiera Ortiz-Severín, Constanza Lecaros, Ian Pérez, Macarena A. Varas, Miguel L. Allende, Francisco P. Chávez

## Abstract

Understanding how bacterial metabolism shapes virulence requires infection models that are both biologically informative and experimentally accessible. Here, we used zebrafish (*Danio rerio*) larvae infection by static immersion as a simple and robust vertebrate model to study *Pseudomonas aeruginosa* host-pathogen interactions and virulence regulation. Using this approach, we show that disruption of polyphosphate (polyP) synthesis by loss of *ppk1* gene (τ1*ppk1*) leads to marked attenuation of virulence, whereas interrupting *ppk2* gene (τ1*ppk2*) results in a hypervirulent phenotype. Phenotypic assays and quantitative proteomics revealed that loss of polyP selectively impairs pyoverdine production, with minimal effects on other canonical virulence factors. Consistently, a pyoverdine-deficient mutant (τ1*pvd*F) exhibited reduced virulence in zebrafish, validating the capacity of the zebrafish model to functionally resolve key virulence factors. Together, our results highlight polyphosphate metabolism as a central regulator of *P. aeruginosa* virulence and position zebrafish immersion assays as an efficient and ethically aligned alternative to mammalian models for studying bacterial pathogenesis.

## Introduction

*Pseudomonas aeruginosa* is a ubiquitous Gram-negative bacterium and a major opportunistic pathogen responsible for acute and chronic infections, particularly in immunocompromised individuals and patients with cystic fibrosis, burns, or implanted medical devices ^1,2^. Its pathogenic success relies on a remarkable capacity to integrate environmental and metabolic cues to coordinate the expression of virulence determinants, including siderophores, phenazines, proteases, and secreted toxins ^3,4^. This tight coupling between metabolism and virulence underlies the adaptability of *P. aeruginosa* across diverse host environments ^5–7^.

Phosphate (Pi) availability is a key metabolic signal that reshapes *P. aeruginosa* physiology and virulence ^8^. Pi limitation activates the PhoB/R regulatory system, which intersects with networks that control iron acquisition, secondary metabolite production, and secreted virulence factors, thereby coupling nutrient stress to pathogenic potential ^9^. Because inorganic polyphosphate functions as an intracellular phosphate reservoir and regulatory hub ^10–12^, alterations in polyP metabolism provide a direct link between Pi-responsive physiology and virulence regulation during infection. PolyP is a highly conserved metabolic polymer involved in energy buffering, metal homeostasis, stress tolerance, and global regulatory control ^13^. In bacteria, polyP is primarily synthesized by polyphosphate kinase 1 (PPK1), and deletion of *ppk1* has been consistently associated with attenuated virulence in multiple pathogens, including *P. aeruginosa* ^14–18^. Despite the robustness of this phenotype, the specific virulence determinants affected by polyP dysregulation and the mechanisms linking polyP metabolism to pathogenic outcomes remain incompletely understood.

In addition, although previous work has identified certain virulence-associated traits linked to disruption of polyphosphate kinase 2 (PPK2), including modest effects on biofilm formation and on secreted factors such as pyoverdine and pyocyanin ^19,20^, the contribution of *ppk2* deletion to overall virulence phenotypes in *P. aeruginosa* remains unclear ^21^. For example, a triple *ppk2A/B/C* mutant exhibits defects in swimming motility and biofilm formation and shows slight reductions in siderophore and phenazine production, yet it maintains near-wild-type levels of total polyP and does not recapitulate the strong attenuation observed in *ppk1* mutants or by complete polyP depletion ^21^. These observations suggest that while PPK2 can influence individual virulence traits, its role in governing virulence outcomes in *P. aeruginosa* during host infection warrants further investigation. Elucidating these metabolic-virulence connections has traditionally relied on mammalian infection models, which, while informative, are constrained by ethical considerations, experimental complexity, and limited throughput. Consequently, there is growing interest in alternative vertebrate models that enable functional interrogation of host-pathogen interactions while adhering to the principles of the 3Rs (Replacement, Reduction, and Refinement) ^22–24^. Zebrafish (*Danio rerio*) larvae provide a powerful system for infection biology ^25^, combining optical transparency, conserved innate immune responses, genetic tractability, and compatibility with scalable experimental designs.

Zebrafish infection studies have commonly employed microinjection-based approaches, which offer precise control over bacterial delivery but require specialized equipment and technical expertise. As an alternative, static immersion assays provide a simple, noninvasive route of infection that more closely resembles natural exposure. We previously established and validated a zebrafish larval immersion protocol as a robust model for studying bacterial pathogenesis and host innate immune responses, demonstrating its ability to resolve virulence phenotypes and inflammatory outcomes *in vivo* ^26–29^. Building on this framework, the present study extends the application of immersion-based infection to dissect the metabolic regulation of virulence in *P. aeruginosa*.

Here, we use static larval infection in zebrafish larvae to investigate the role of polyphosphate metabolism in *P. aeruginosa* pathogenicity. We show that loss-of-function mutation of *ppk1* (Δ*ppk1*) results in a pronounced attenuation of virulence *in vivo*, associated with a selective impairment in pyoverdine production. Functional validation using a pyoverdine-deficient mutant confirms the central role of this virulence factor in zebrafish infection. In addition, our data reveal previously uncharacterized effects of the Δ*ppk2* mutation, underscoring the need for further investigation of its role in infection biology.

Together, our findings establish polyP metabolism as a critical regulatory layer linking bacterial physiology to virulence and demonstrate that zebrafish larvae provide an efficient, ethically aligned platform for dissecting host-pathogen interactions and identifying key pathogenic determinants in *P. aeruginosa*.

## Results

### Polyphosphate mutants exhibit altered virulence in zebrafish and differential expression of virulence factors

To investigate the contribution of polyphosphate metabolism to virulence, we performed a series of *in vivo* infection assays and *in vitro* phenotypic tests using zebrafish larvae and phosphate-regulated bacterial cultures. Zebrafish larvae at 3 days post-fertilization were statically immersed in suspensions of *P. aeruginosa* PAO1 wild-type (WT), Δ*ppk1*, or Δ*ppk2* mutants grown in either low-phosphate (↓Pi) or high-phosphate (↑Pi) conditions.

Under ↓Pi conditions, wild-type PAO1 induced over 70% mortality in zebrafish larvae within 24 hours, consistent with its known increased virulence in phosphate-limiting environments. In contrast, larvae exposed to the Δ*ppk1* mutant displayed survival rates above 80% at 48 hours post-infection, regardless of phosphate condition. This attenuation closely matched the survival observed in larvae infected with the avirulent *E. coli* DH5α control strain, and no significant differences were observed between *E. coli* DH5α and the Pa Δ*ppk1* mutant at ↑Pi conditions (Figure 1A). However, *E. coli* DH5α did not cause any physiological alterations (or symptoms) in the larvae throughout the course of the infection, whereas Δ*ppk1* mutants affected larvae over time, increasing infection severity in 22% of larvae at 24 hpi (Figure 1B). At his time, 42% and 62% of larvae infected with Pa WT in ↑Pi and ↓Pi, respectively, showed an increased infection severity. These data confirm that mutation of *ppk1* significantly impairs the bacterium’s virulence and lethality.

**Figure 1.**
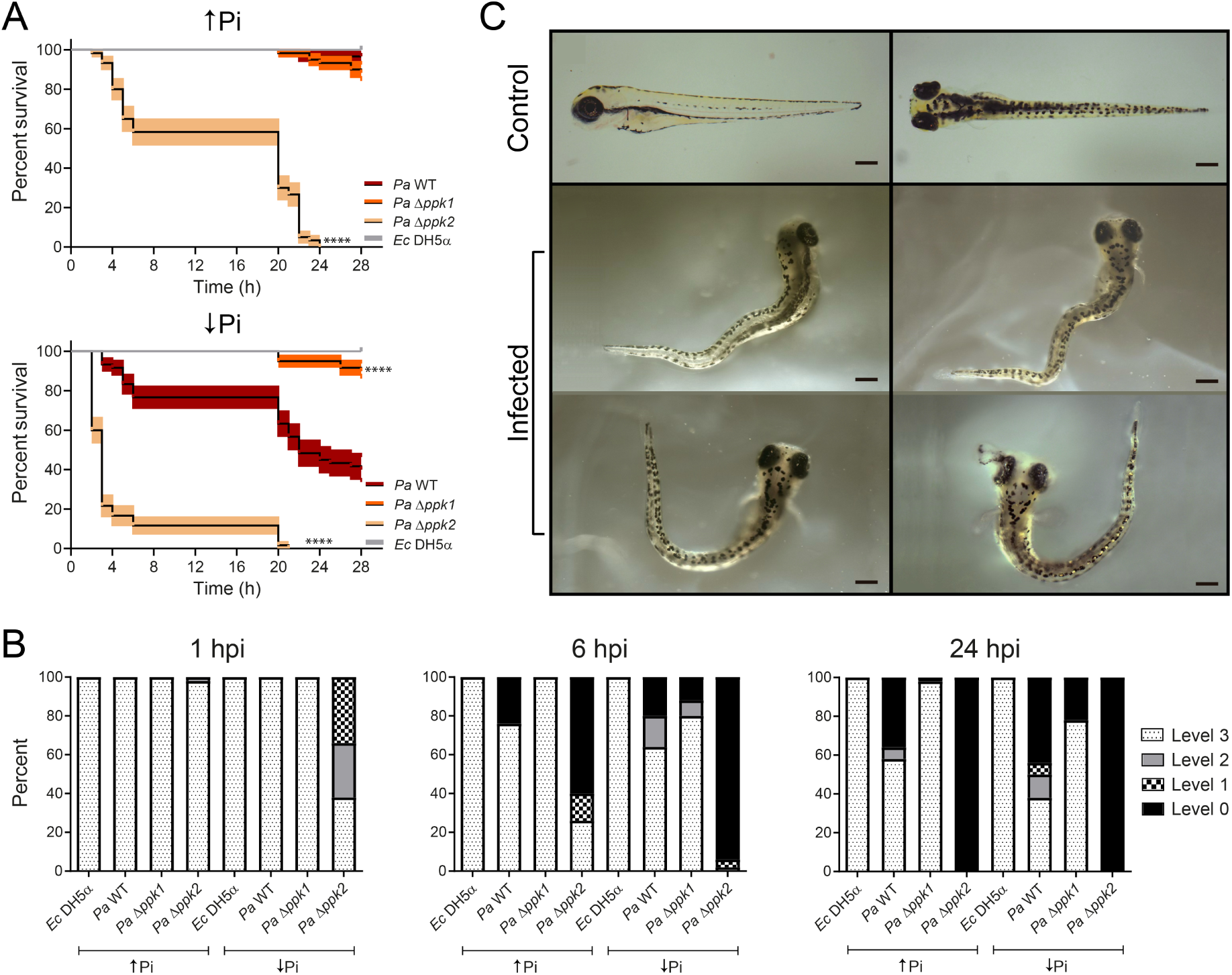
Effect of zebrafish infection by immersion with *P. aeruginosa* WT and polyphosphate mutants’ strains. Zebrafish larvae (3 dpf) exposed to different *P. aeruginosa* strains by static immersion. Bacteria were grown to the stationary state in broth media supplemented with phosphate (↑Pi) or without phosphate (↓Pi), and about 2×10^8^ CFU/mL were suspended in sterile fish media (E3) for the immersion experiments. (A) Survival curves. Asterisks denote significant differences between the polyP synthesis mutants and the WT strain (Logrank (Mantel-Cox) test Holm-Šídák’s multiple comparisons post-test, ****p < 0.0001). *E. coli* DH5α was used as a control. (B) Pathological progression of the infected larvae. The graph shows the percent of larvae presenting symptoms corresponding to one of 4 infection severity levels: Level 3 = no clinical signs, normal larvae; Level 2= reduced response to touch, decreased blood flow in tail; Level 1= decreased heart rate and/or pericardial edema, unresponsive to touch; Level 0 = death. (C) Representative images of zebrafish larvae after *P. aeruginosa* infection by immersion and their external symptoms. Bar = 100 µm.

Strikingly, the Δ*ppk2* mutant exhibited 100% larval mortality within 24 hours post-infection under both ↓Pi and ↑Pi conditions. This result was further supported by survival curve analysis, which showed a sharp decline in the viability of larvae infected with the Δ*ppk2* mutant, compared with the more gradual decline observed with *Pa* WT strain. In addition, larvae showed increased infection severity as soon as 1 hpi (Figure 1B), suggesting a strongly hypervirulent phenotype. Infected zebrafish larvae displayed marked phenotypic alterations compared to controls, including a progressive deterioration of physiological and behavioral status. Finally, as shown in Figure 1C, larvae exhibited pronounced body curvature, loss of normal straight posture, and clear signs of paralysis, indicating severe neuromuscular impairment. Notably, several larvae also showed ocular infection, as previously reported in *P. aeruginosa* mammalian infections ^30,31^. These phenotypes correlate with the progression of infection severity described in Figure 1B, where larvae transition from asymptomatic states (Level 3) to reduced touch response, decreased blood flow, pericardial edema, and ultimately death (Level 0).

To identify possible mechanisms behind these phenotypes, we assessed the production of key secreted virulence factors on specific indicator media. Interestingly, rhamnolipid, elastase, protease, and hemolysin production were the same for the polyP synthesis mutants and the WT strain, irrespective of Pi availability (except for elastase, which was moderately increased in the Δ*ppk1* mutant in ↑Pi, Figure 2). Only siderophore production was significantly increased in the Δ*ppk2* mutant, both in ↑Pi and ↓Pi (Figure 2), suggesting that siderophores are the principal contributors to the observed virulence shift.

**Figure 2.**
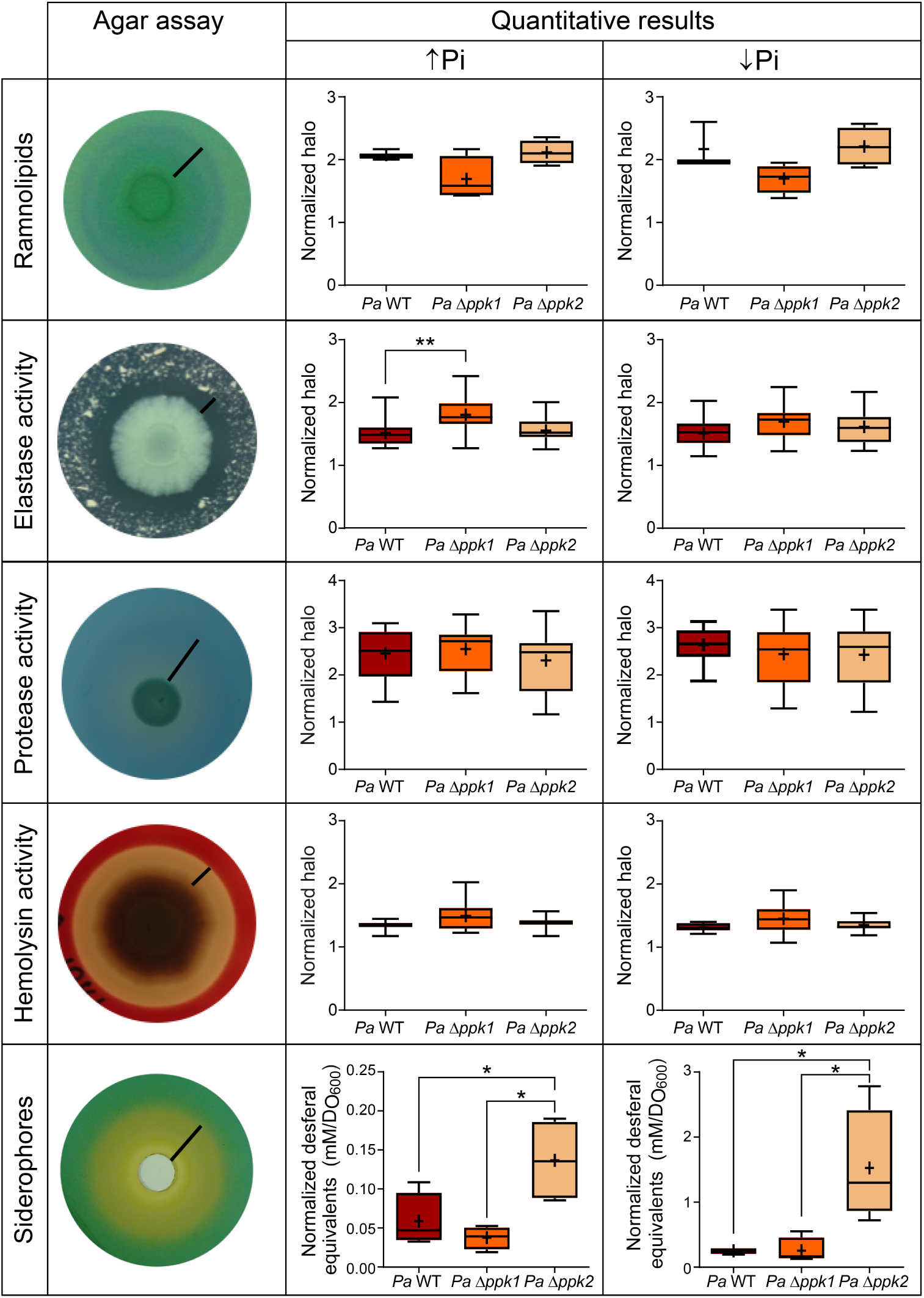
Virulence factor production in the polyP synthesis mutants. Production of virulence factors was evaluated using indicator agars, as shown in the left panel. The black bar indicates the halos produced by the bacterial virulence factors, which were measured and plotted (graphs in the central and right panels). Box plots of quantified virulence factors. The halos were normalized to the optical density (OD_600_) of the cultures used as the assay inoculum. For siderophore production, desferal equivalents were calculated from a calibration curve and then normalized to the OD_600_ of the cultures used in the assay. Asterisks denote significant differences between the polyP synthesis mutants and the WT strain (One-way ANOVA with Tukey’s multiple comparison post-test, **p < 0.01, *p < 0.05).

Together, these results demonstrate that PPK1 and PPK2 exert opposing regulatory effects on *P. aeruginosa* virulence and that their influence is closely linked to the regulation of iron-scavenging and redox-active virulence factors.

### Quantitative proteomics reveals proteome-wide effects of polyphosphate metabolism

To elucidate the molecular basis underlying the divergent virulence phenotypes observed in zebrafish, we performed quantitative proteomic analyses using ICPL labeling under phosphate-limited conditions. Differential expression of proteins was observed in the WT strain in response to Pi limitation, and also between the WT and the polyP synthesis mutants in ↓Pi (Table 1). Comparison of the WT strain grown under low versus high phosphate identified 125 differentially expressed proteins, confirming extensive metabolic and regulatory remodeling in response to phosphate availability (Figure 3A).

**Figure 3.**
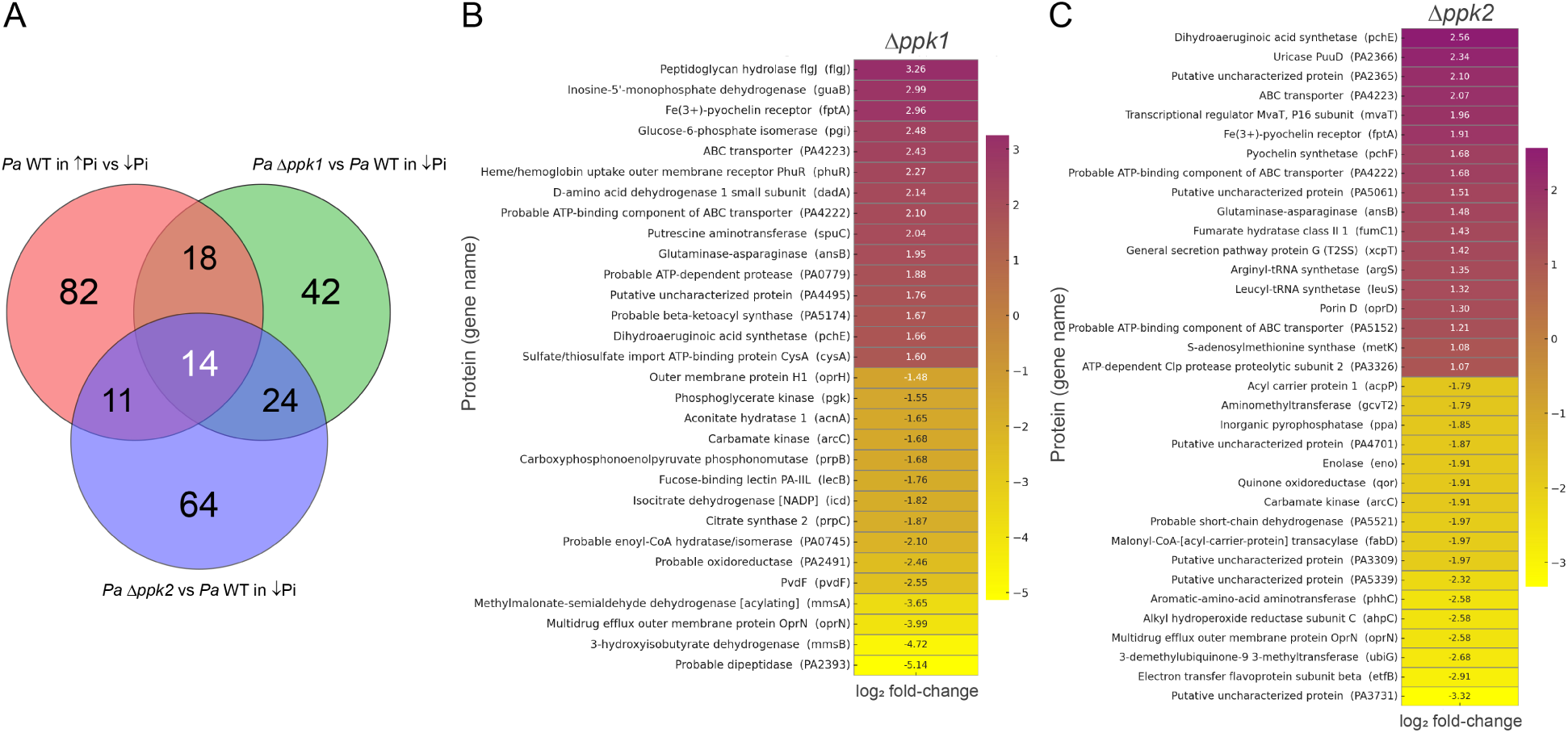
Quantitative proteomic analysis of polyphosphate kinase mutants in *Pseudomonas aeruginosa*. (A) Venn diagram showing differentially abundant proteins in *P. aeruginosa* WT strain under low versus high phosphate (↓Pi vs ↑Pi) and in Δ*ppk1* and Δ*ppk2* mutants relative to WT under phosphate-limited conditions. (B) Heatmap of the top 15 differentially abundant proteins in the Δ*ppk1* mutant, highlighting reduced abundance of pyoverdine biosynthesis proteins. (C) Heatmap of the top 15 differentially abundant proteins in the Δ*ppk2* mutant, showing increased abundance of proteins associated with phenazine metabolism and transport functions. Proteins are ranked by absolute log₂ fold change (|log₂FC| > 1).

**Table 1.** Summary of quantitative proteomics comparisons under phosphate-limited conditions.

| Comparison | Total peptides | Quantified peptides | Identified proteins | Changed proteins |
| --- | --- | --- | --- | --- |
| *Pa* WT ↓Pi vs *Pa* WT ↑Pi | 3602 | 2054 | 414 | 125 |
| *Pa Δppk1* ↓Pi vs *Pa* WT ↓Pi | 2816 | 1818 | 372 | 98 |
| *Pa Δppk2* ↓Pi vs *Pa* WT ↓Pi | 5096 | 2833 | 528 | 113 |

Disruption of *ppk1* resulted in a more selective proteomic reprogramming, with 98 proteins differentially abundant relative to the WT strain under low-phosphate conditions, 42 of which were unique to the Δ*ppk1* mutant (Figure 3A). Notably, several enzymes directly involved in pyoverdine biosynthesis were significantly downregulated, including PvdF, PvdA, and PvdH, along with additional proteins associated with iron acquisition and oxidative stress management (Figure 3B). Proteins involved in phosphate uptake and metabolism, as well as envelope-associated functions, were also reduced in abundance.

In contrast, the Δ*ppk2* mutant displayed a broader proteomic response, with 113 differentially expressed proteins, 64 of which were unique to this mutant (Figure 3A). Among the proteins showing increased abundance were enzymes involved in phenazine biosynthesis and modification, including PhzA1, PhzB1, PhzM, and PhzS, as well as proteins involved in iron metabolism and redox processes (Figure 3C). In addition, elevated levels of outer membrane and transport-associated proteins, including components of the MexEF-OprN efflux system and porins such as OprD, were detected.

Both polyphosphate kinase mutants exhibited altered abundance of proteins associated with envelope remodeling and global regulation (Figure 3). Increased levels of the transcriptional regulator MvaT were detected in both Δ*ppk1* and Δ*ppk2* backgrounds, suggesting that perturbation of polyP metabolism impacts higher-order regulatory networks rather than isolated virulence factors.

Considering that pyoverdine and phenazine biosynthesis and modification pathways were differentially expressed in the polyP synthesis mutants, and that they are important virulence factors relevant for *P. aeruginosa* infection, we evaluated and compared their production (Figure 4). Parallel *in vitro* analyses revealed that the Δ*ppk1* mutant exhibited impaired pyoverdine production, whereas pyocyanin levels varied with phosphate conditions. The Δ*ppk2* mutant produced elevated levels of both pyoverdine and pyocyanin. These findings suggest that polyphosphate metabolism modulates virulence by regulating siderophore and phenazine production, with opposing effects depending on the polyP kinase deleted. The targeted proteomic signature closely mirrored the pronounced defect in pyoverdine production and the attenuated virulence phenotype observed in zebrafish for the Δ*ppk1* mutant. For the *ppk2* mutant, the observed proteomic changes were consistent with the increased production of pyocyanin and pyoverdine observed phenotypically, although they do not define a single dominant virulence pathway.

**Figure 4.**
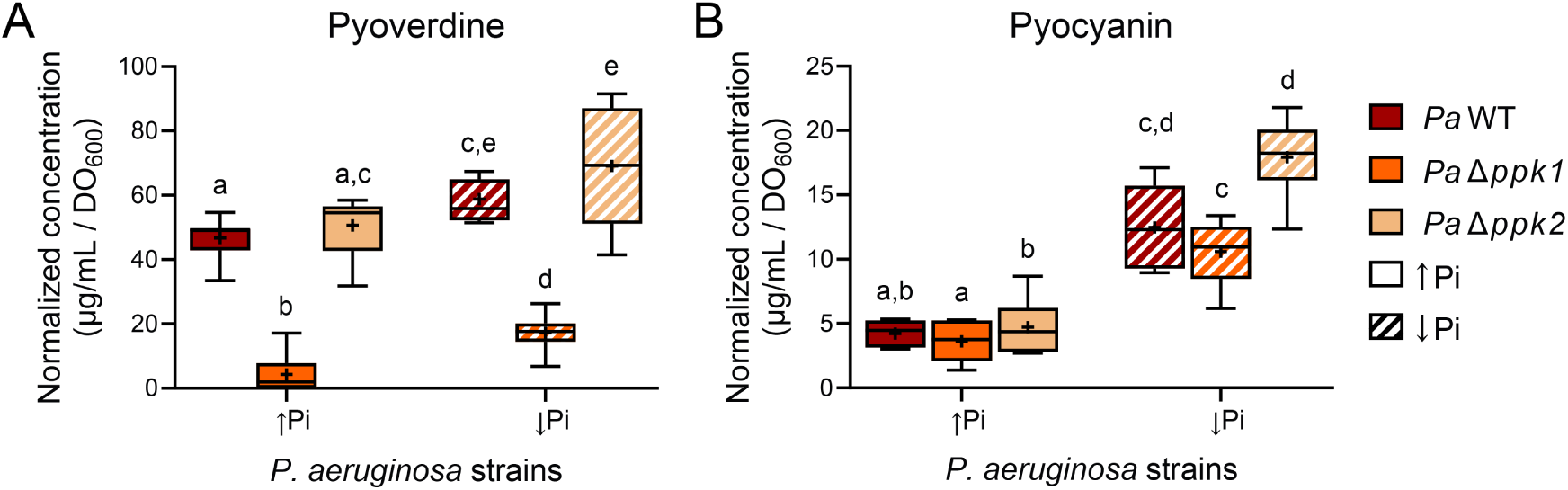
Quantification of relevant virulence factors identified by proteomics. Pyoverdine (A) and pyocyanin (B) quantification based on absorbance methods using a standard curve. Box plots show the average (“+” symbol) values (±SD) of at least three independent replicates. Different letters above the boxes indicate significant differences between the Δ*ppk1* or Δ*ppk2* mutant and the WT strain across Pi conditions (Two-way ANOVA with Tukey’s multiple comparisons post hoc test).

Together, these data indicate that PPK1-dependent polyphosphate metabolism sustains a specific virulence program centered on pyoverdine biosynthesis, whereas disruption of *ppk2* leads to a more diffuse proteomic remodeling involving phenazine production, transport systems, and envelope-associated functions. This distinction provides a molecular framework for understanding the attenuated versus hypervirulent phenotypes observed *in vivo*.

### Pyoverdine production is required for full virulence in zebrafish

To functionally validate the contribution of pyoverdine to *P. aeruginosa* virulence, zebrafish larvae were infected with a pyoverdine-deficient mutant (Δ*pvdF*) and compared to the WT strain using the immersion assay. Loss of pyoverdine biosynthesis resulted in a pronounced attenuation of virulence, with significantly increased larval survival under both phosphate-replete and phosphate-limited conditions (Figure 5A). This phenotype closely mirrored the reduced lethality observed for the Δ*ppk1* mutant, supporting a mechanistic link between polyphosphate metabolism, pyoverdine production, and *in vivo* pathogenicity.

**Figure 5.**
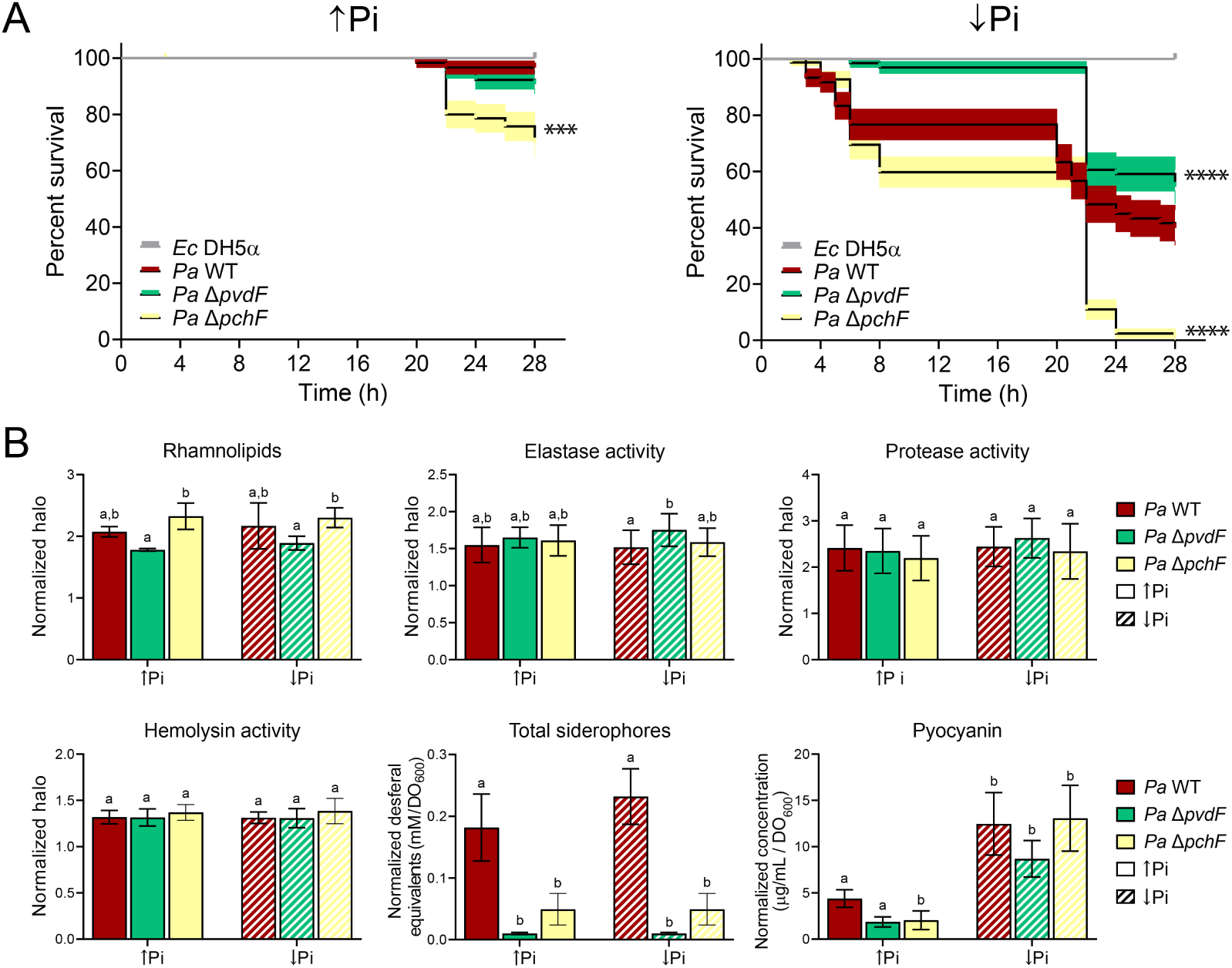
Survival curves and virulence factor production of mutants defective in siderophore synthesis. (A) Survival curves. Asterisks denote significant differences between the polyP synthesis mutants and the WT strain (Logrank (Mantel-Cox) test Holm-Šídák’s multiple comparisons post-test, ***p < 0.001, ****p < 0.0001). *E. coli* DH5α was used as a control. (B) Virulence factor production. Different letters above the boxes indicate significant differences between the polyP synthesis and the WT strain at different Pi conditions (Two-way ANOVA with Tukey’s multiple comparisons post-test).

Consistent with this outcome, quantitative analyses confirmed that the Δ*pvdF* mutant was severely impaired in total siderophore production, while levels of other secreted virulence factors, including elastase, protease, rhamnolipids, hemolysin, and pyocyanin, were largely comparable to those of the wild-type strain (Figure 5B). Interestingly, a significant increase in elastase activity was observed only in the Δ*pvdF* mutant (in ↓Pi conditions), similar to what occurred in the Δ*ppk1* mutant (Figures 2 and 5). These results indicate that the observed attenuation cannot be attributed to a generalized defect in virulence factor expression, but rather to the specific loss of pyoverdine-dependent functions.

To assess the possibility that the impaired lethality of the Δ*ppk1* mutant was linked to decreased production of other siderophores besides pyoverdine, we evaluated the survival of zebrafish larvae infected with a pyochelin-deficient strain (Δ*pchF* mutant). Unexpectedly, the Δ*pchF* mutant exhibited higher lethality in the zebrafish immersion model than the Δ*ppk1* mutant and the WT strain (Figure 5A). This was not explained by an increase in the production of other virulence factors (Figure 5B), so the global effect of pyochelin production during *P. aeruginosa* infection should be further inspected. Together, these data establish pyoverdine as a key determinant of *P. aeruginosa* virulence in the zebrafish model and provide functional validation of the proteomic signature observed in the Δ*ppk1* mutant.

### Pyocyanin contributes to virulence-associated traits but is not sufficient to drive lethality

Given the increased production of pyocyanin observed in the Δ*ppk2* mutant and its association with hypervirulence, we next evaluated the role of pyocyanin biosynthesis in zebrafish infection using mutants defective in phenazine production (Δ*phzM*, Δ*phzS*, and Δ*phzR*). In contrast to the strong attenuation observed for the pyoverdine-deficient strain, disruption of pyocyanin synthesis resulted in non-significant differences in zebrafish survival when compared to the WT strain, irrespective of Pi conditions (Figure 6A). None of the pyocyanin-deficient mutants phenocopied the avirulent behavior of Δ*ppk1* or Δ*pvdF*, indicating that loss of pyocyanin alone is insufficient to abolish virulence in this model.

**Figure 6.**
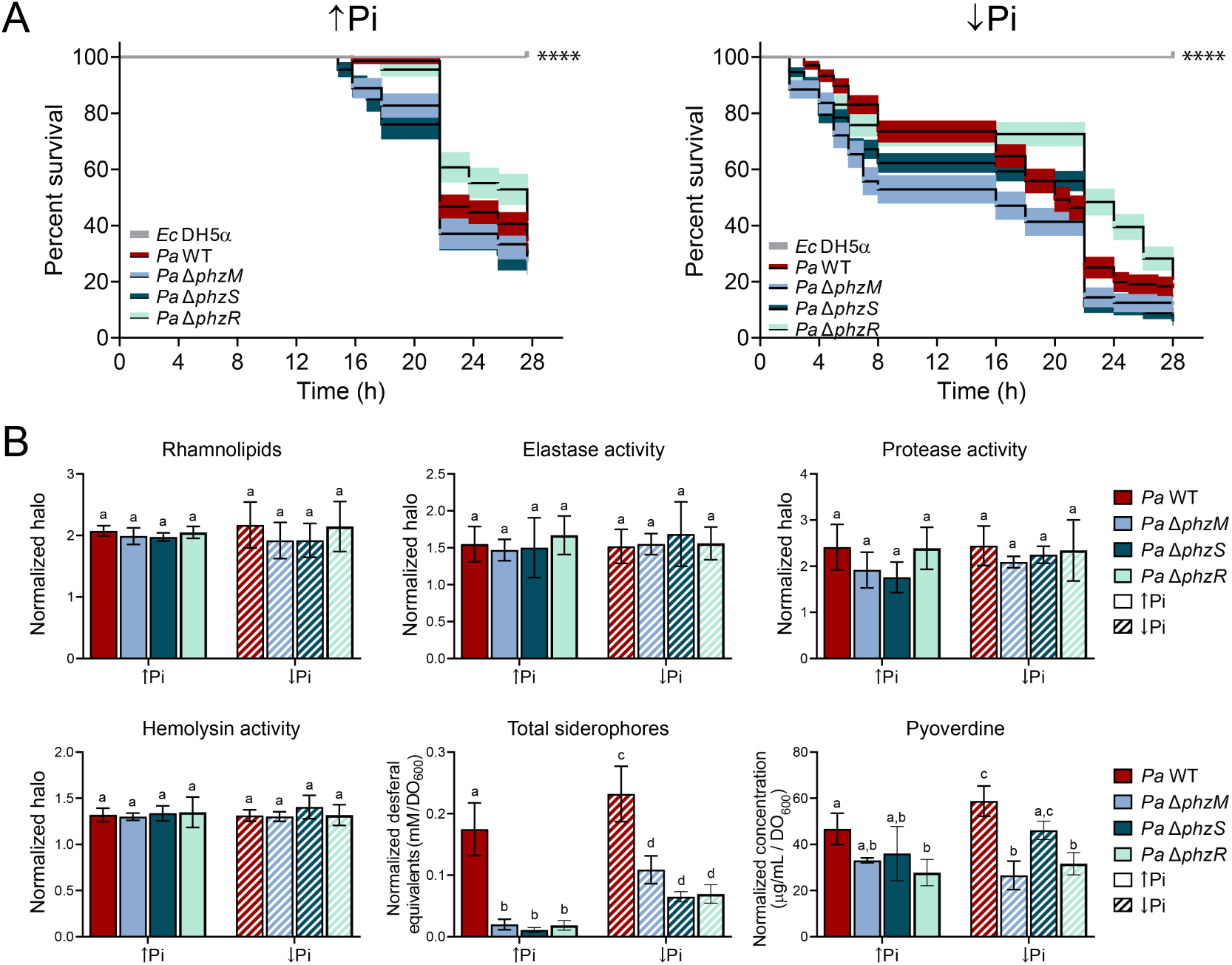
Survival curves and virulence factor production of mutants defective in pyocyanin synthesis. (A) Survival curves. Asterisks denote significant differences between the polyP synthesis mutants and the WT strain (Logrank (Mantel-Cox) test Holm-Šídák’s multiple comparisons post-test, ***p < 0.001, ****p < 0.0001). *E. coli* DH5α was used as a control. (B) Virulence factor production. Different letters above the boxes indicate significant differences between the polyP synthesis mutants and the WT strain at different Pi conditions (Two-way ANOVA with Tukey’s multiple comparisons post-test).

Phenotypic characterization confirmed that pyocyanin-deficient mutants retained normal or near-normal production of other virulence factors, including rhamnolipids, elastase, proteases, and hemolysins (Figure 6B). Notably, pyoverdine levels were mainly preserved in these backgrounds, except for Δ*phzM* in ↓Pi and Δ*phzR* in both Pi conditions, highlighting a complex interconnection between iron acquisition and redox-active toxin production as the dominant driver of lethality in zebrafish larvae under the conditions tested.

Together, these findings demonstrate that while pyocyanin contributes to the virulence-associated phenotype of *P. aeruginosa*, particularly in the context of altered polyphosphate metabolism, it does not represent a primary determinant of host lethality in zebrafish. Instead, our data position pyoverdine-mediated iron acquisition as the central effector linking polyphosphate metabolism to virulence outcomes in this vertebrate infection model.

## Discussion

In this study, we used a zebrafish larval infection model to examine how polyphosphate metabolism modulates *Pseudomonas aeruginosa* virulence *in vivo*. Our results show that deletion of *ppk1* leads to a marked attenuation of virulence, correlating with a strong reduction in pyoverdine production, whereas deletion of *ppk2* results in a hypervirulent phenotype associated with increased levels of both pyoverdine and pyocyanin. These contrasting outcomes highlight the distinct and nonredundant roles of polyphosphate kinases in coordinating bacterial metabolic state with pathogenic behavior during host infection.

The central role of pyoverdine in the attenuated phenotype of the Δ*ppk1* mutant is consistent with extensive evidence showing that iron acquisition systems are critical determinants of *P. aeruginosa* virulence ^20,32^. Phosphate limitation is known to activate regulatory networks that link phosphate sensing to iron uptake, redox balance, and virulence factor secretion^33,34^. Within this framework, polyphosphate functions as an intracellular phosphate reservoir and regulatory integrator, allowing bacteria to translate external nutrient cues into coordinated physiological responses. Our data extend this model by demonstrating that disruption of polyphosphate synthesis selectively impairs siderophore production and compromises virulence in a vertebrate host^35^.

Proteomic analyses further support this interpretation by revealing targeted downregulation of pyoverdine biosynthetic proteins in the Δ*ppk1* mutant, rather than a global collapse of virulence-associated functions. This pattern indicates that polyphosphate metabolism fine-tunes specific virulence pathways that are particularly relevant under *in vivo* conditions. In contrast, the broader proteomic remodeling observed in the Δ*ppk2* mutant, including increased abundance of proteins associated with phenazine biosynthesis, iron-related functions, and transport systems, suggests a different mode of regulatory perturbation. The contribution of PPK2 to virulence appears more context dependent, consistent with its proposed role in modulating nucleotide and redox metabolism rather than directly controlling siderophore biosynthesis.

Phosphate availability has emerged as a host-relevant signal during infection, influencing virulence expression across multiple experimental systems. Nutrient limitation can promote pathogenic programs that enhance iron scavenging, oxidative stress resistance, and host tissue damage. Our results show that these phosphate-responsive virulence traits are recapitulated in zebrafish larvae and critically depend on intact polyphosphate metabolism, positioning polyP as a central intracellular mediator linking phosphate sensing to virulence regulation.

The zebrafish larval model provides important advantages for investigating these processes. As a non-mammalian vertebrate host, zebrafish combine conserved innate immune responses with experimental scalability and optical accessibility, enabling functional dissection of host–pathogen interactions while adhering to the principles of Replacement, Reduction, and Refinement. The ability of simple, immersion-based infections to resolve virulence differences driven by metabolic regulation underscores the utility of this system for studying nutrient-dependent pathogenic mechanisms relevant to human infection.

Taken together, our findings identify polyphosphate metabolism, and specifically PPK1-dependent pathways, as key modulators of the phosphate-iron-virulence axis in *P. aeruginosa*. By integrating *in vivo* infection assays, quantitative proteomics, and targeted genetic analyses, this work provides a mechanistic link between bacterial metabolic regulation and pathogenic outcomes. More broadly, it establishes zebrafish larvae as a robust and ethically aligned platform for investigating virulence regulation in opportunistic human pathogens, offering a complementary approach to mammalian infection models.

## Conclusion

This study demonstrates that inorganic polyphosphate metabolism is a key regulator of *Pseudomonas aeruginosa* virulence, with distinct contributions from PPK1- and PPK2-dependent pathways. Disruption of *ppk1* resulted in a pronounced attenuation of virulence, which was closely associated with impaired pyoverdine production, whereas alteration of *ppk2* led to a hypervirulent phenotype, accompanied by increased expression of iron- and redox-related virulence factors.

Using zebrafish larvae as a vertebrate infection model, we show that simple immersion-based assays can reliably distinguish attenuated from hypervirulent bacterial phenotypes and elucidate the *in vivo* contributions of specific virulence determinants. Quantitative proteomic analyses provided mechanistic support for these observations by revealing targeted remodeling of virulence-associated pathways rather than global disruption of the proteome. Together, our findings highlight polyP metabolism as an important regulatory layer linking bacterial metabolic state to pathogenic outcomes and establish zebrafish larvae as a robust and ethically aligned platform for studying host-pathogen interactions. The integration of *in vivo* infection models with proteomic profiling offers a general framework for dissecting virulence mechanisms in opportunistic human pathogens while reducing reliance on mammalian models.

## Materials and Methods

### Bacterial strains and cell conditions

*P. aeruginosa* PAO1 wild-type (WT) and mutant strains were obtained from the *Pseudomonas* Genome Database collection ^36^. All *P. aeruginosa* strains were routinely grown in LB broth or LB agar. Phosphate-replete (↑Pi) and phosphate-limited (↓Pi) conditions were created by using a modified PGS culture medium (3g/L NaCl, 2% glycerol, 2.5 g/L peptone, 1 mM CaCl_2_ and 1m MgSO_4_), supplemented with 2.5% v/v of KPh buffer at pH = 6 for the ↑Pi condition. For phosphate-regulated experiments, overnight cultures of *P. aeruginosa* strains in LB were washed twice with the corresponding ↑Pi or ↓Pi culture media and used as inoculum for 250 mL flasks (1:100 dilution). Cultures were incubated for 18 h at 37 °C and 180 rpm, and optical density was measured at 600 nm (OD_600_), with cultures adjusted with fresh medium as needed.

### Zebrafish husbandry

Zebrafish (*Danio rerio*) embryos were obtained by natural spawning of Tab5 (wild type) line^37^. Fertilized eggs were raised in Petri dishes containing E3 medium (5 mM NaCl, 0.17 mM KCl, 0.33 mM CaCl_2_, 0.3 mM MgSO_4_) and 0.1% methylene blue until 3 days post fertilization (dpf). All procedures complied with national guidelines of the Animal Use Ethics Committee of the University of Chile and the Bioethics Advisory Committee of Fondecyt-Conicyt.

### Bacterial immersion and injection experiments

The bacterial cultures were washed and subsequently suspended for immersion assays in filter-sterilized system water (60 mg/L Instant Ocean Sea Salts (Aquarium Systems, Mentor, OH, USA; pH 7.0). The cell suspension was adjusted to OD_600_ = 1.4. The bacterial suspension was serially diluted and plated on LB-agar to determine the number of viable cells (CFU/mL). 6-dpf zebrafish were washed with system water, and 10 larvae were placed per well in a sterile 6-well plate. The wells were filled with the bacterial solution to obtain an OD_600_ of 0.5, equivalent to 5×10^8^ CFU/mL in a final volume of 8 mL. The zebrafish were incubated at 28°C until 3 dpf, and then the temperature was changed to 22°C prior to the co-incubation with bacteria. The larvae were placed in 6-well plates containing bacterial suspensions, and the progression of infection was monitored up to 28 hours post-infection (hpi). Kaplan-Meier survival analyses were performed to evaluate zebrafish lethality in bacterial immersion and injection assays.

### Virulence assays in indicator agars

Indicator agars were used to quantify rhamnolipid (CTAB-methylene agar)^38^, elastase (elastin-nutrient plate^39^, protease (milk agar, Biomerieux) and hemolysin (Sheep-blood agar, Biomerieux) activity. In each indicator agar, 5 µL of *P. aeruginosa* WT or mutant strain cultures grown in either ↑Pi or low ↓Pi, were inoculated on top of the agar. Plates were incubated at 37°C for 24 hours before measuring the halo.

Total siderophore production (CAS agar plate without nutrients) was quantified by measuring the orange halo produced by sterile supernatants (50 µL each) of grown cultures of *P. aeruginosa* strains placed in holes made in CAS plates^40^. A calibration curve was made using different concentrations of desferal (deferoxamine mesylate, Calbiochem, Merck) ranging from 0.1 mM to 1 mM desferal diluted in ↑Pi and ↓Pi media. Plates with desferal or bacterial supernatant were incubated at room temperature for 48 hours before measuring the halo. Desferal-equivalents were calculated based on the equations obtained from the calibration curves per each medium.

For all virulence agars, graphs were made by dividing the measured halos (or desferal equivalents) by the OD_600_ of the culture used to inoculate the indicator agar. For all experiments, at least three independent replicates were made.

### Pyoverdine and pyocyanin quantification

The production of pyoverdine and pyocyanin in *P. aeruginosa* WT and mutant strains was quantified by measuring the absorbance of culture supernatants in a spectrophotometer. First, a spectrophotometric scan between 200 and 500 nm of pyoverdine and pyocyanin pure compounds (Sigma-Aldrich) was made to establish the optimum absorbance for each compound in the different culture media (pyoverdine: 383 nm in ↑Pi and 386 in ↓Pi; pyocyanin: 311 nm in ↑Pi and ↓Pi). After, calibration curves were created by measuring the absorbance of solutions of pyoverdine from 0-100 µg/mL, and pyocyanin from 0-20 µg/mL in ↑Pi and ↓Pi media. Calibration curves were used to calculate pyoverdine or pyocyanin equivalents from spent supernatants, which were then normalized to the OD_600_ of their corresponding cultures.

### Quantitative proteomics

Quantitative proteomic analyses were performed using isotope-coded protein labeling (ICPL) to compare *Pseudomonas aeruginosa* PAO1 WT and polyphosphate mutants grown under phosphate-replete (↑Pi) and phosphate-limited (↓Pi) conditions. Total protein extracts were obtained from three independent biological replicates per condition, pooled to generate representative samples (150 μg total protein per condition), quantified using the Bio-Rad Protein Assay, and stored at −20 °C prior to analysis.

For ICPL labeling, 100 μg of each protein sample was solubilized in 8 M urea and 25 mM ammonium bicarbonate, reduced and alkylated, then diluted to 2 M urea and digested overnight with trypsin (20:1, w/w) at 37 °C. Peptides were desalted using C18 tips and resuspended in 0.2% trifluoroacetic acid. Labeling was performed at the peptide level using ICPL reagents according to the manufacturer’s instructions and previous analysis ^41^. Briefly, peptides were incubated with ICPL reagent in guanidinium chloride buffer for 2 h at 25 °C under an inert atmosphere, and reactions were quenched with hydroxylamine. Differentially labeled samples were then combined, chemically neutralized, dried, and stored at −20 °C.

Combined ICPL-labeled samples (∼200 μg) were fractionated using an offline high-pH reversed-phase chromatography, generating 25–30 fractions per experiment. Each fraction was subsequently analyzed by nanoLC-ESI-MS/MS using an Ultimate 3000 nanoHPLC system coupled to an HCT Ultra ion trap mass spectrometer (Bruker Daltonics). Peptides were separated on a C18 reversed-phase column under a linear acetonitrile gradient in 0.1% formic acid at a flow rate of 300 nL/min. Mass spectrometry data were acquired in positive ion mode using data-dependent acquisition, collecting full MS scans (m/z 350–1500) followed by MS/MS fragmentation of the four most abundant ions, with dynamic exclusion applied.

Raw MS/MS data from all fractions were merged and processed using DataAnalysis (Bruker Daltonics), and peptide identification was performed using Mascot against a *P. aeruginosa* reference proteome database supplemented with reversed sequences for false discovery rate (FDR) estimation. Search parameters included carbamidomethylation (fixed), methionine oxidation and ICPL labeling (variable), and allowance of one missed cleavage. Peptide identifications were filtered to an FDR ≤ 5%, and only proteins identified with at least two peptides were considered for further analysis.

Quantitative analysis was performed using WARP-LC software (Bruker Daltonics), based on extracted ion chromatograms of ICPL-labeled peptide pairs ^42,43^. Relative protein abundances were calculated from monoisotopic peak intensities, log₂-transformed, and median-normalized. This approach enabled robust comparative analysis of proteome remodeling associated with phosphate availability and polyphosphate metabolism.

## Acknowledgments

We are grateful to Nicole Molina for technical assistance at SysmicroLab. This work was partly supported by FONDECYT grants 1221360 (FCH and MA) and 1262424 (FCH) and. CONICYT fellowships supported MV (3170449) and JOS (21130717).

